# On information metrics for spatial coding

**DOI:** 10.1101/189084

**Authors:** Bryan C. Souza, Rodrigo Pavão, Hindiael Belchior, Adriano B.L. Tort

## Abstract

The hippocampal formation is involved in navigation, and its neuronal activity exhibits a variety of spatial correlates (e.g., place cells, grid cells). The quantification of the information encoded by spikes has been standard procedure to identify which cells have spatial correlates. For place cells, most of the established metrics derive from Shannon’s mutual information (Shannon, 1948), and convey information rate in bits/sec or bits/spike (Skaggs et al., 1993; Skaggs et al., 1996). Despite their widespread use, the performance of these metrics in relation to the original mutual information metric has never been investigated. In this work, using simulated and real data, we find that the current information metrics correlate less with the accuracy of spatial decoding than the original mutual information metric. We also find that the top informative cells may differ among metrics, and show a surrogate-based normalization that yields comparable spatial information estimates. Since different information metrics may identify different neuronal populations, we discuss current and alternative definitions of spatially informative cells, which affect the metric choice.

## Introduction

The hippocampus is known to be involved in memory formation (Scoville and Milner, 1957; Eichenbaum, 2000) and spatial navigation (O’Keefe and Dostrovsky, 1971; Morris et al., 1982; Zola-Morgan and Squire, 1990; O’Keefe and Recce, 1993). In the early 70s, O’Keefe and Dostrovsky discovered that some hippocampal cells have firing rate modulated by the animal’s position, discharging more at a spatial region known as the place field of the cell (O’Keefe and Dostrovsky, 1971). Since the discovery of place cells, other types of spatial correlates emerged in related areas to the hippocampal circuitry, such as the head-direction cells in the postsubiculum (Taube et al., 1990), and the grid cells and speed cells in the entorhinal cortex (Fyhn et al., 2004; Hafting et al., 2005; Moser et al., 2008; Kropff et al., 2015). Properly identifying these cells requires estimates of the information contained in spikes about navigational features (i.e., position, speed, head angle). The main metrics used to estimate this type of information were proposed by Skaggs et al. (1993, 1996) and are derivations from Shannon’s mutual information (MI).

Information entropy, as originally proposed by Shannon, measures the amount of uncertainty in the outcome of a variable based on its probability of occurrence (Shannon, 1948). In other words, the more unpredictable the outcome is, the more entropy it has. On the other hand, the MI is a measure of the shared entropy between two variables; it indicates how much knowing a variable X reduces the uncertainty of a variable Y. While the MI is usually measured in bits, the two derived metrics proposed by Skaggs et al. express information in bits per spike (I_spike_) or bits per second (I_sec_). These metrics are not straightforward divisions of the MI by the number of spikes or elapsed time, but are rather defined as estimates of the average information rate conveyed by the cell (Skaggs et al., 1993); this is achieved by keeping the lower order terms of the MI power series expansion with respect to time (see Methods). A fundamental difference from the MI is that I_sec_ and I_spike_ only take into account the average firing rate over the spatial variable (i.e., location, speed), ignoring firing rate differences across multiple occurrences of the same variable (e.g., across trials). Although these metrics provide a meaningful interpretation of the relation between firing rate and navigational features, the possible implications introduced by these modifications remain to be investigated. Of note, the MI has been previously used to measure spatial information (Ego-Stengel and Wilson, 2007), though a direct comparison with the I_sec_ and I_spike_ metrics has never been performed.

In this work, we use simulated data to address how well the metrics I_spike_, I_sec_ and MI reflect the capacity of decoding the animal’s position based on the spikes of an individual neuron, which directly relates to the amount of spatial information conveyed by the cell (Quiroga and Panzeri, 2009). We find that while MI values correlate well with decoding performance under a variety of scenarios, this is not always the case for I_spike_ and I_sec_. Similar results hold when analyzing real spikes from place cells of rats recorded on a linear track. Moreover, we also find that the different metrics may give rise to different subpopulations of spatially modulated neurons. Finally, we show that a surrogate-based normalization can equalize the three metrics. We end by discussing the conceptual definition of spatially informative cells, which may vary according to the employed metric.

## Methods

### Simulating spatially modulated cells

We simulated the firing rate of spatially modulated neurons across 30 trials on a linear track divided into 25 bins of space (Figure 1A). For simplicity, the animal speed and occupancy were considered constant over space. We modeled eight types of cells; for each cell type, we simulated 10 levels of spatial modulation (Figure 1B, Neuron ID a to j). For the first 5 cell types (Neuronal type I to V), the firing rate of each trial was modeled as a Gaussian centered (on average) at bin 13 with (average) standard deviation of 5 bins. To introduce inter-trial variability, 0.5 and 0.1 white noise was added to the center and standard deviation of the Gaussian, respectively. The cell types mimicked the behavior of (I) a pyramidal-like place cell (low basal firing rate); (II) an interneuron-like place cell (high basal firing rate); (III) interneuron-like place cell negatively modulated by space; (IV) a pyramidal-like place cell exhibiting spatial modulation in a subset of trials; and (V) a cell as in IV, but with constant mean firing rate across trials. For cell types I-III, we varied spatial modulation strength (deviation from baseline), while for cell types IV and V the number of modulated trials varied. Cell type VI was similar to cell type I but could have multiple, equally-spaced peaks as a grid cell. We also simulated cells behaving as ramp (VII) or constant functions (VIII) along space, with different slopes and firing rate levels, respectively.

**Figure 1.**
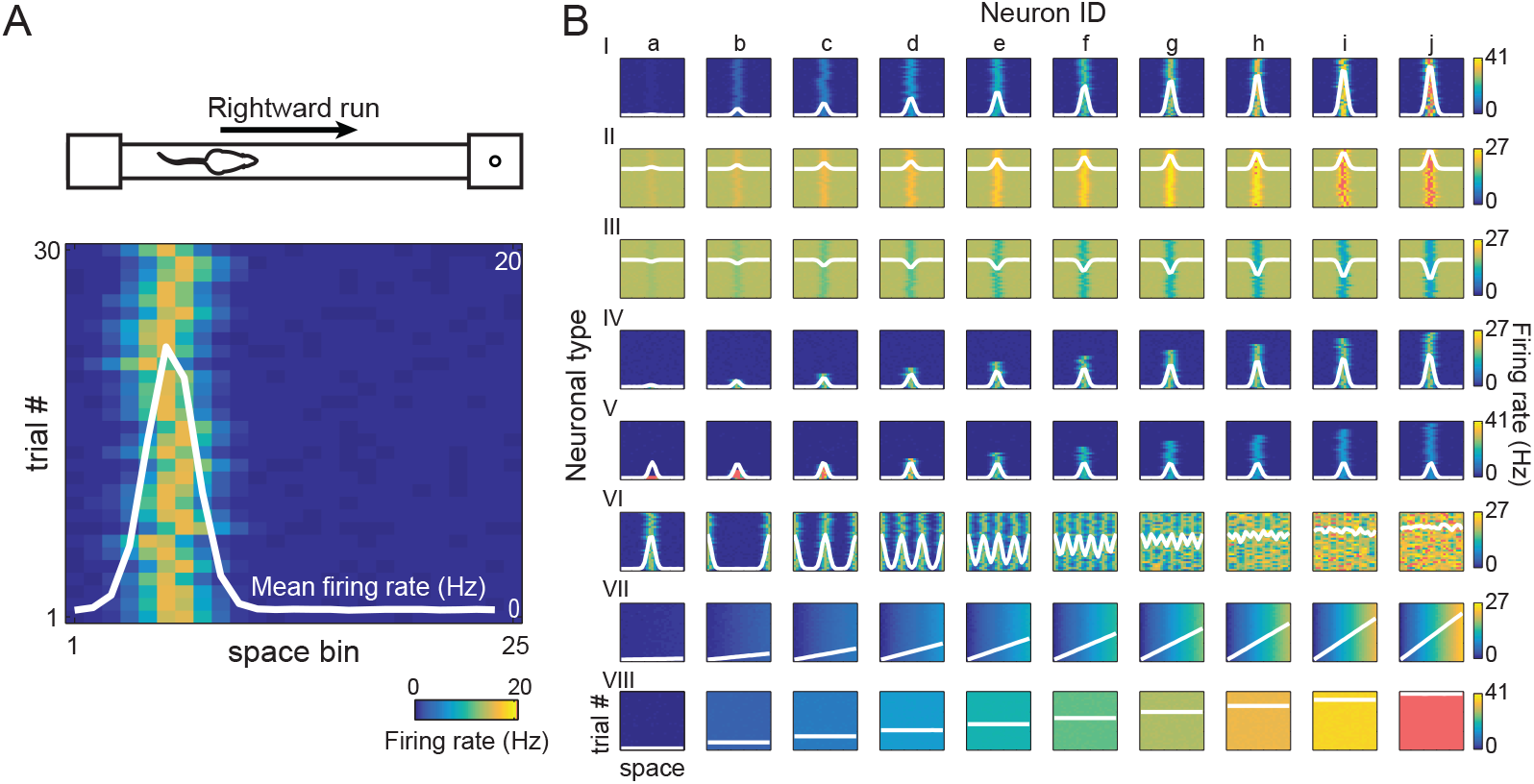
Simulating the activity of spatially modulated neurons. **A.** Neuronal firing rate was simulated in 25 bins of space over 30 trials, meant to represent spiking activity of one cell during rightward runs on a linear track. In the rate-position map, pseudo-colors represent firing rate on each trial, while the white line is the mean firing rate over trials. **B**. Rows display 8 simulated neuronal types (I-VIII) of different spatial modulation profiles. Cell types I-V exhibit place-cell-like behavior, while types VI, VII and VIII represent grid cells, ramp and constant functions, respectively (see Methods for details). For each type, columns show the rate-position maps (pseudocolor scale) as well as the mean firing rate (overlaying white trace) for 10 distinct cells (a-j) differing in the level of spatial modulation.

### Estimating spatial information in the firing rate

To estimate the spatial information contained in the firing rate of each cell, we used I_spike_ and I_sec_ – the main metrics for selecting place cells (Skaggs et al., 1993, 1996) ‒ and the MI (Shannon, 1948). We computed the I_sec_ metric from the average firing rate (over trials) in the 25 space bins using the following definition:

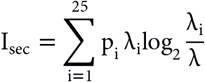

where λ_i_ is the mean firing rate in the i-th space bin and p_i_, the occupan cy ratio of the bin, while λ is the overall mean firing rate of the cell. I_sec_ measures information rate in bits per second (see next section for mathematical derivation).

The I_spike_ metric is a normalization of I_sec_, defined as:

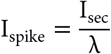

This normalization yields values in bits per spike. The MI was estimated using all firing rate values (within trials), which were binned into four non-overlapping quantiles:

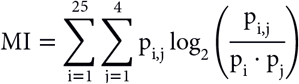

where p_i_ and p_i_ are the probabilities of position bin *i* and firing rate bin *j*, respectively; p_ij_ is the joint probability between position bin *i* and firing rate bin *j*.

### Derivation of I_sec_ from MI

Using Bayes rule, the MI can be rewritten as:

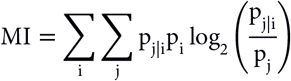

where p_j|i_ is the conditional probability of firing rate bin *j* given the position *i*. For a sufficient small amount of time Δt, we can assume that a cell can either emit just one spike or none, and thus there are only two possible firing rate bins, denoted as *j*=0 (no spike) and *j*=1 (spike). The probability of spike occurrence during Δt at position *i* is given by p_j=1|i_ = λ_i_Δt, while p_j=0|i_ = 1 - λ_i_Δt denotes the probability of no spike. Similarly, *p*_j=1_ = λΔt and p_j=1_ = 1 - λΔt denote spiking probabilities irrespective of position. Using these probabilities in the MI formula above gives:

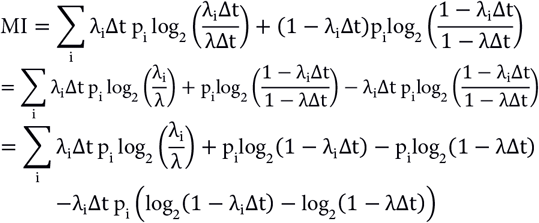

Using the first term in the power series expansion of logarithms, we have that:

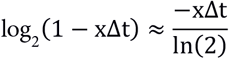

Applying this approximation in the previous MI equation yields:

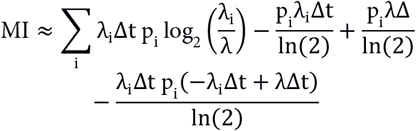

Excluding the second order terms (Δt^2^), we have:

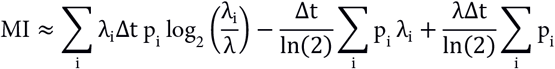

Using that λ = Σ_i_p_i_ λ_i_ and Σ_i_p_i_ = 1, the last two terms cancel out, yielding:

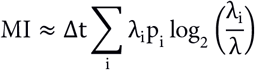

Finally, since the power series expansion of MI around t is given by:

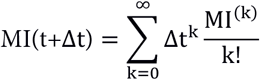

where MI^(k)^ is the k-th time derivative, we have that the first time derivative is approximated by:

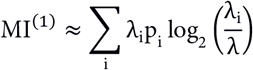

which is the definition of I_sec_.

### Estimating spatial information from decoding performance

Decoding algorithms predict the most likely stimulus that generated a given response based on the previous observations of stimulus-response pairs (Quiroga and Panzeri, 2009). We used a Gaussian naïve Bayes classifier to predict the position of the animal based on the firing rate of a cell (see John and Langley, 1995 for detailed description). This approach defines a conditional probability model with the prior probability of position and the likelihood probability of firing rate given the position, which is assumed to be normally distributed. These probabilities are estimated based on the available samples and used to compute the posterior probability of position given the firing rate. New firing rate samples can then be assigned to the most probable position as defined by the *maximum posterior probability*.

We performed the classification using a leave-one-out approach (Figure 2). Briefly, all firing rate values (across positions and trials; training bins) but one (test bin) are used to estimate the posterior probabilities (Figure 2A). The model then predicts the position of the left out firing rate value. This procedure is repeated multiple times so that each firing rate value is used once as a test bin. A confusion matrix is next constructed from actual and decoded positions, and used to extract the percentage of correct decoding, which is the proportion of entries in the y=x diagonal (Figure 2B). In this work, the percentage of correct decoding was assumed to correlate with the true spatial information content of the cell.

**Figure 2.**
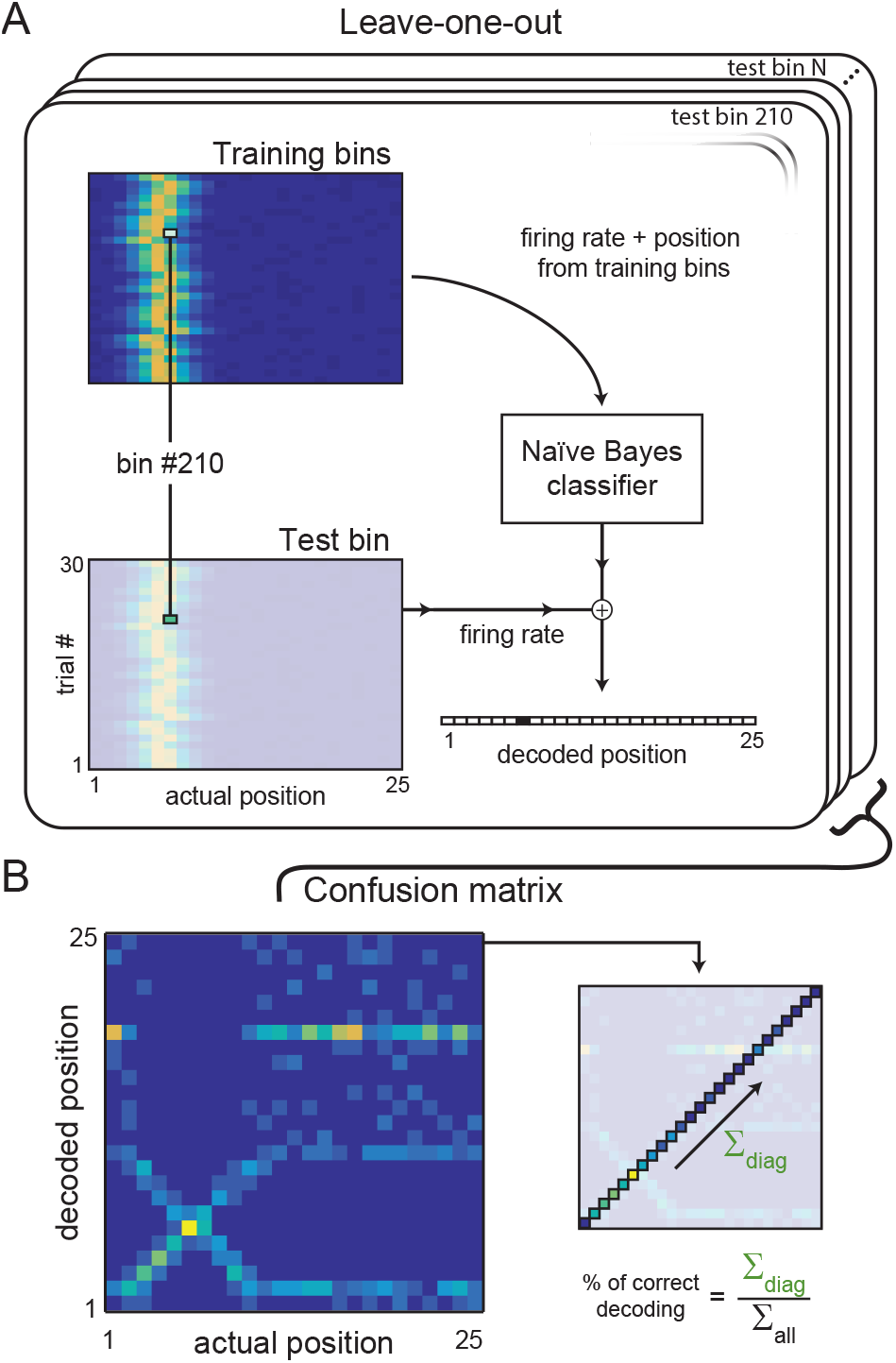
Estimating spatial information using a Bayesian classifier. **A.** All the (training) bins of the rate-position map except one were used to train a naïve Bayes classifier. The classifier was then used to decode the position of the remaining (test) bin using only its firing rate. Each bin was used once as a test bin (leave-one-out approach). **B.** We used all the decoded positions to compute the confusion matrix, which relates decoded and actual positions. We divided the amount of times the classifier was correct (sum of the matrix diagonal) by the total number of classifications to calculate the decoding performance.

### Normalizing spatial information metrics using surrogates

To normalize the estimates of spatial information, we first shuffled the labeling of position bins on each trial (Figure 4A). This approach avoids any firing preference across trials and in the mean rate. We then computed the I_spike_, I_sec_ and MI metrics using the shuffled bins. This procedure was repeated 100 times and used to build a surrogate distribution for each metric. The information values were then expressed as z-scores of the surrogate distribution. The z-scored metrics are referred to as normalized I_spike_ (Norm. I_spike_), I_sec_ (Norm. I_sec_) and MI (Norm. MI).

### Measuring spatial information in real neurons

To investigate the relation between the spatial information estimates and decoding performance for real cells, we used a dataset with recordings from the CA1 region of the dorsal hippocampus of three rats running back and forth on a linear track (Mizuseki et al., 2013; data freely available at <u>https://crcns.org/</u>). We calculated the spatial information estimates and the decoding performance for all neurons in 75 recording sessions. In Figure 8, we used the classification of putative interneurons and pyramidal cells available in the dataset (Mizuseki et al., 2009).

### Comparing the subpopulations of spatially-informative neurons

We investigated the overlap between the subpopulations of most informative neurons according to the different metrics. First, for each metric we ranked the cells from most to least informative. We then computed the percentage of common cells between each pairwise combination of metrics for the N% most informative cells, with N% varying from 20% to 100%.

## Results

We first computed the three spatial metrics for each of the 8 simulated cell types shown in Figure 1. These cell types vary in how their firing rate is modulated by space (see Methods). For instance, while cell type I fires at its place field location on every trial, cell type IV emits spatially modulated spikes only in a subset of trials. For each cell type, we varied the amount of spatial modulation in 10 levels. This was achieved by either changing the deviation of the firing rate from the basal level (cell types I-III), or the percentage of trials with modulated activity (cell types IV and V), number of place fields (cell type VI), spatial slope (cell type VII) or basal firing rate level (cell type VIII). The spatial metrics were then compared with the percentage of correct decoding of a Bayesian classifier.

Figure 3A,B shows examples in which the same decoding performance could have either high or low values of I_sec_ and I_spike_ (compare different rows in Figure 3B). Moreover, there were also cases in which I_sec_ and I_spike_ were insensitive to changes in the percentage of correct decoding (see cell type V in Figure 3B). Across all the simulated cells, we found a clear correlation between the percentage of correct decoding in log scale and MI, but not between correct decoding and either I_sec_ or I_spike_ (Figure 3C).

**Figure 3.**
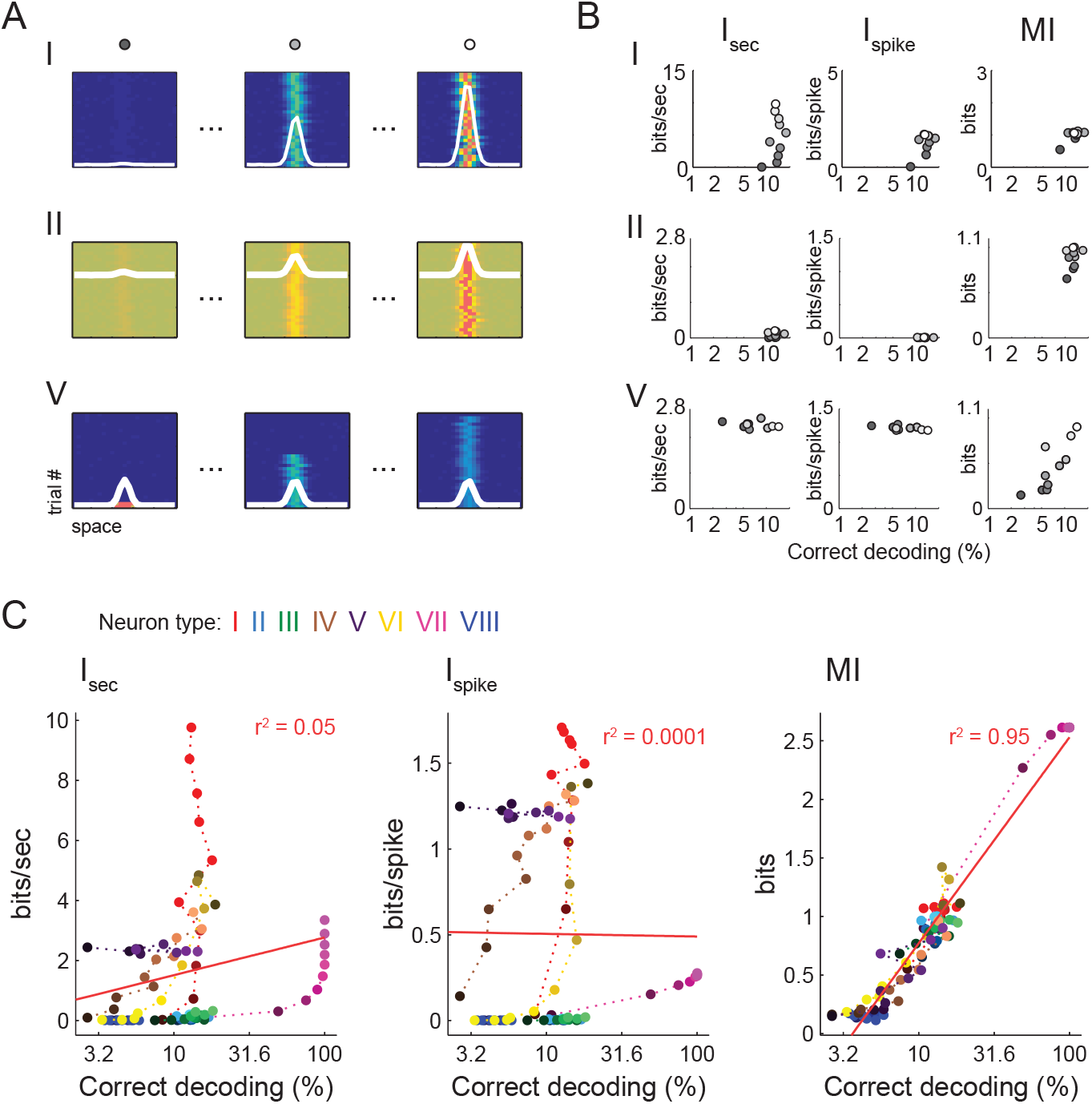
Assessing the performance of spatial information metrics. **A.** Examples of cell types I, II and V during three modulation conditions. Circles on top indicate the strength of spatial modulation, from black (low) to white (high). **B.** I_sec_, I_spike_ and MI values of the cell types in A plotted against the percentage of correct decoding of the animal’s position (see Methods). Note that the same level of decoding across different cell types can elicit distinct I_sec_ and I_spike_. values (i.e., compare rows), and that similar values of I_sec_ or I_spike_ within a same cell type may be associated to different decoding levels (i.e., cell type V). **C.** Percentage of correct decoding in log scale for all simulated cells vs. I_sec_, I_spike_ and MI along with the linear fit (red line). Colors denote neuronal types. Dotted line and color gradient go from the lowest (dark) to highest (light) modulation levels (Neuron ID in Figure 1B). The MI best correlates with decoding performance.

We next corrected the spatial information metrics for the chance information level of each cell, which varies according to firing characteristics. To that end, we computed information in shuffled rate-position maps to build a surrogate distribution of information values (Figure 4A). The actual value was then compared to and z-scored in relation to the chance distribution. While the correlation between decoding performance and MI did not considerably change, the correlation of I_sec_ and I_spike_ with decoding substantially improved after the normalization and became similar across metrics (Figure 4B).

**Figure 4.**
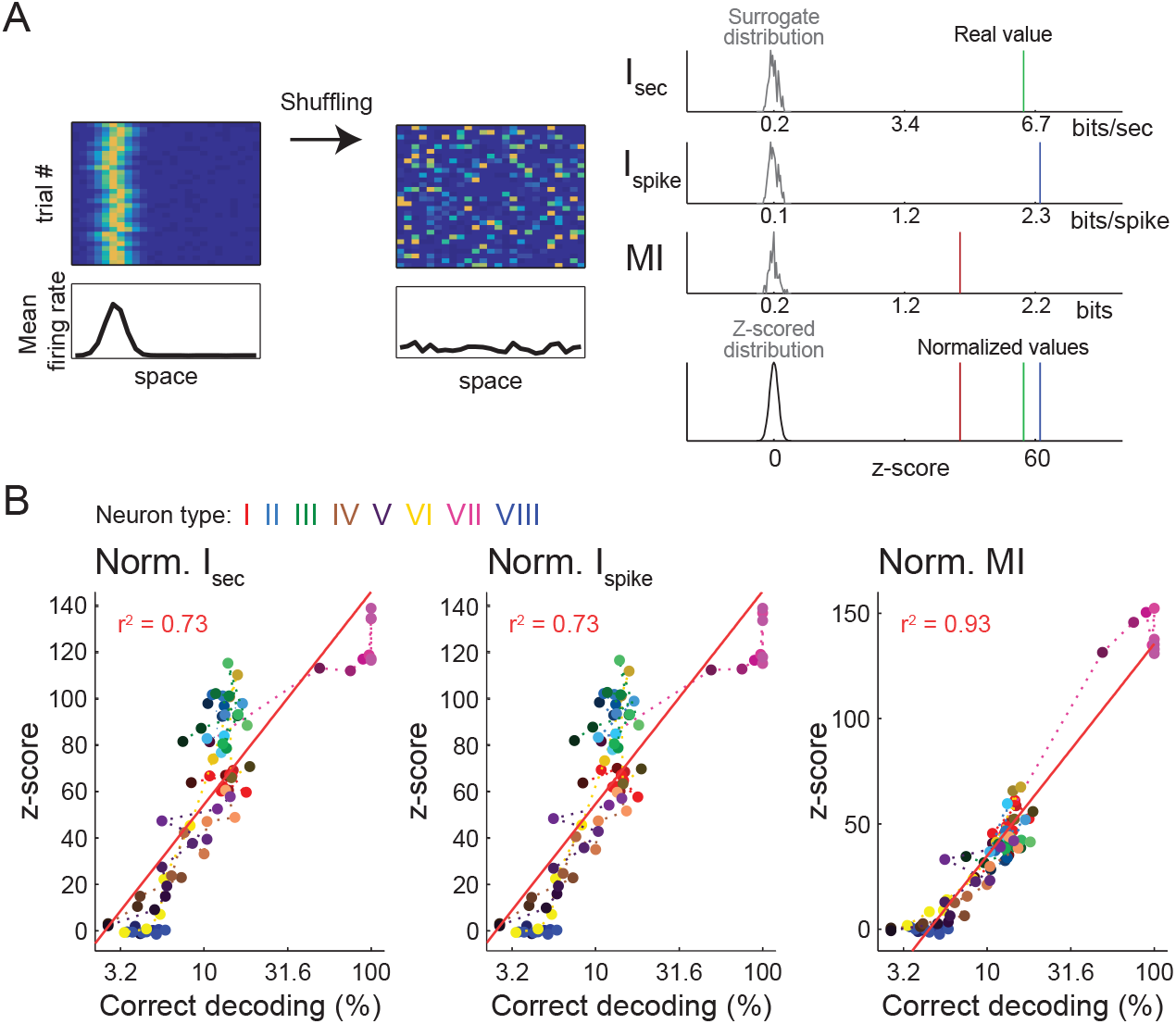
Correcting spatial metrics using surrogates. **A.** (Left) Example of shuffling procedure. To estimate chance information levels, the firing rate bins within trials were shuffled prior to computing the metrics. (Right) The actual value of each metric was compared to the surrogate distribution (n=100 shufflings) and z-score normalized. **B.** Percentage of correct decoding (in log scale) vs. the normalized values of I_sec_, I_spike_ and MI for all simulated cells along with the linear fit (red line). Colors denote neuronal types (as in Figure 3). Normalizing I_sec_ and I_spike_ significantly improves their correlation with decoding performance.

We next investigated spikes from real cells recorded from the rat hippocampus during traversals on a linear track. The original and normalized MI, I_sec_ and I_spike_ were computed for each cell, and plotted against decoding performance in log scale. Figure 5 shows results for an example session. Notice in Figure 5B that the MI values correlated well with decoding performance, while I_spike_ showed no clear correlation (MI: P=0.67; I_spike_: r^2^=0.007). Unexpectedly, the I_sec_ values were better correlated to decoding than in our simulations (I_sec_: r^2^=0.51). The correlation with decoding substantially improved for I_sec_ and I_spike_ after the normalization, and reached similar values across all three metrics (Figure 5C). Figure 6A shows the r^2^ values between each metric and decoding performance for 75 linear track sessions. Consistent with the example in Figure 5, we found that MI held the best correlations with decoding performance, followed by I_sec_, while I_spike_ was poorly correlated. After normalization, the three metrics correlated similarly well with decoding. Figure 6B shows the changes in r^2^ between the original and normalized versions of each metric. While the *ŕ* of I_sec_ and I_spike_ significantly increased, the r^2^ changes for MI were not considerably different.

**Figure 5.**
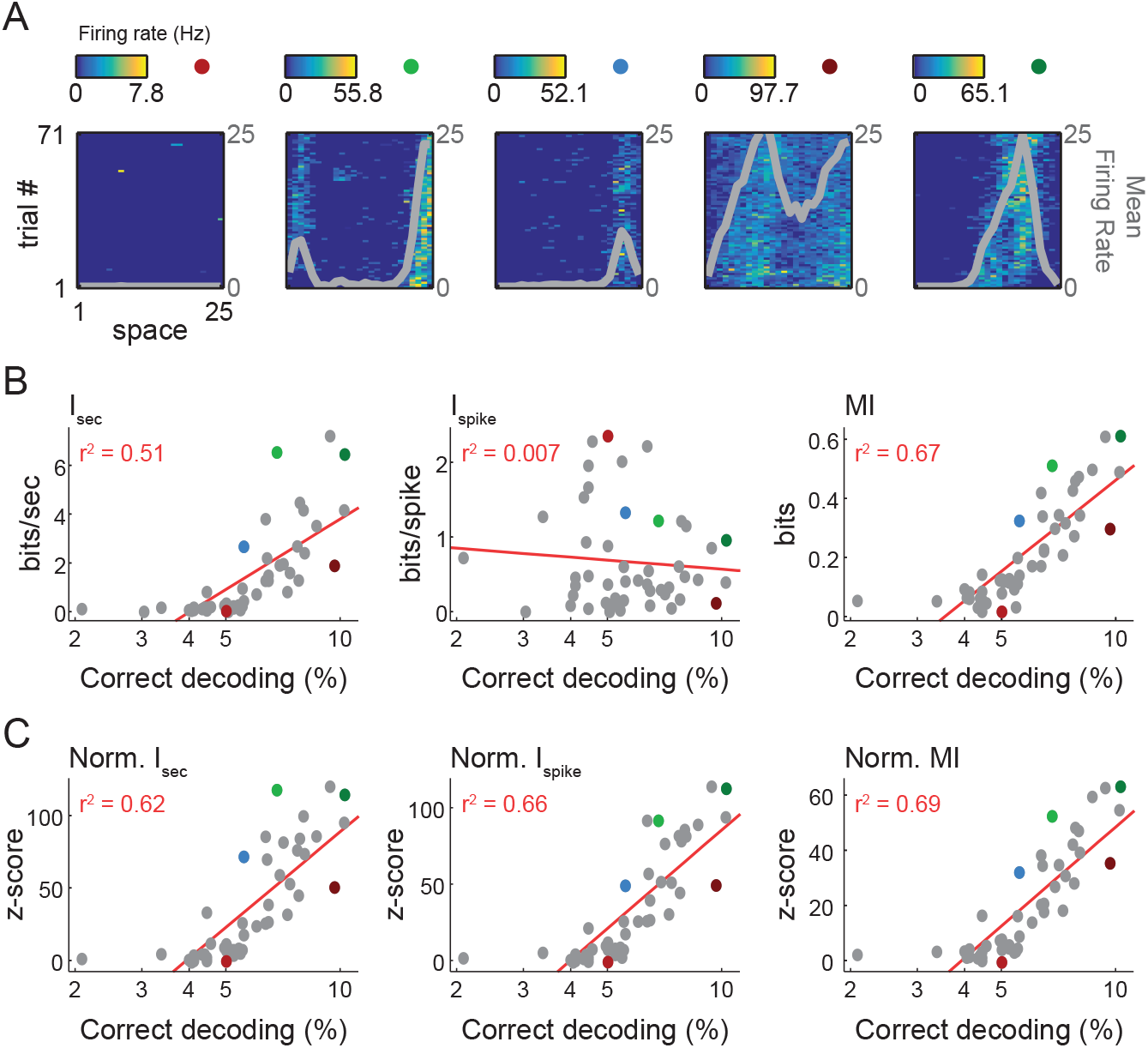
Spatial information metrics applied to real cell data. **A.** Example of the spatial activity of five neurons recorded in hippocampal CA1 of one rat during a linear track session. The recording included spatially modulated (i.e., place cells) as well as nonmodulated cells. Colored circles mark the same cells highlighted in B and C. **B.** Scatter plots of information estimates and decoding performance in log scale for all recorded cells. Note that the MI exhibits the best correlation with decoding performance. **C.** Same as in B but for the normalized metrics. As in simulated data (Figure 3), normalizing I_sec_ and I_spike_ improves their correlation with decoding performance.

**Figure 6.**
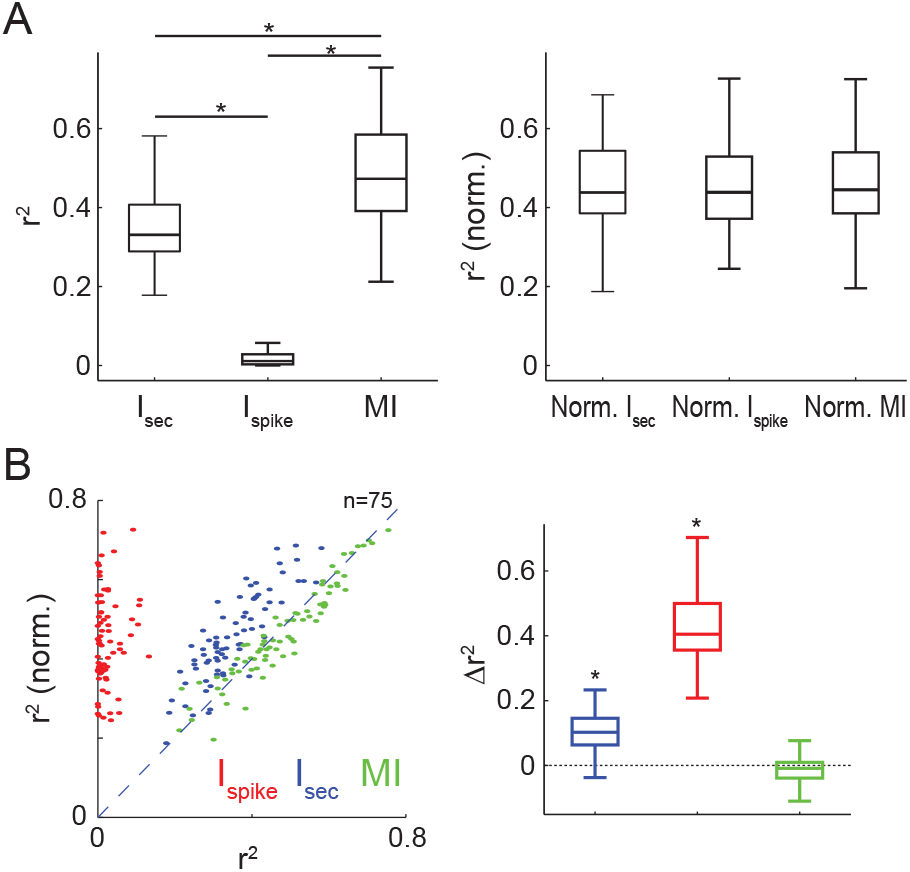
Correlation of spatial metrics and decoding performance. **A.** Boxplots show the distribution of coefficients of determination (r^2^) between the logarithm of the percentage of correct decoding and I_sec_, I_spike_ or MI for 75 linear track sessions. The original and normalized metrics are shown in the left and right panels, respectively. Notice similar r^2^ values for the normalized metrics. *p<0.001 (Wilcoxon signed-rank tests, Bonferroni corrected). **B**. Scatter plot of the r^2^ for each metric and its normalized version (left) and boxplots of the changes in r^2^ after normalization (right). There is a significant increase in r^2^ for I_sec_ and I_spike_; *p<0.001 (Wilcoxon signed-rank test, Bonferroni corrected).

We proceeded to investigate how similar are the subpopulations of spatially-selective cells when defined by each metric at varying degrees of information threshold. To that end, for each metric we ranked neurons according to their spatial information (Figure 7A), and then computed the pairwise intersection (i.e., MI & I_sec_, MI & I_spike_, I_sec_ & I_spike_) between the top N% informative cells. Figure 7B shows the percentage of common cells as a function of N%. For up to the top 40% spatially informative cells, we found that there is very low intersection for I_spike_ and MI and for I_spike_ and I_sec_ (median below 20% of common cells). As expected, the intersection between metrics gradually increases with decreasing the information threshold (i.e., increasing N%), eventually reaching a complete intersection when all cells are analyzed. On the other hand, MI and I_sec_ had a high number of common cells for all information thresholds, with a median intersection of ~90% of cells. Therefore, the subpopulation of spatially informative neurons according to the MI is slightly different from I_sec_ and drastically different from I_spike_ (Figure 7B). Surprisingly, I_sec_ and I_spike_ showed very low overlap among their top informative cells despite their related definitions. We also computed the rank correlation between the metrics and found similar results: a high correlation between MI and I_sec_, and low correlation of I_spike_ with either MI or I_sec_ metrics (Figure 7C left). Interestingly, correlation values increase after normalizing the metrics (Figure 7C right), indicating that the standardized metrics capture similar spatial features.

**Figure 7.**
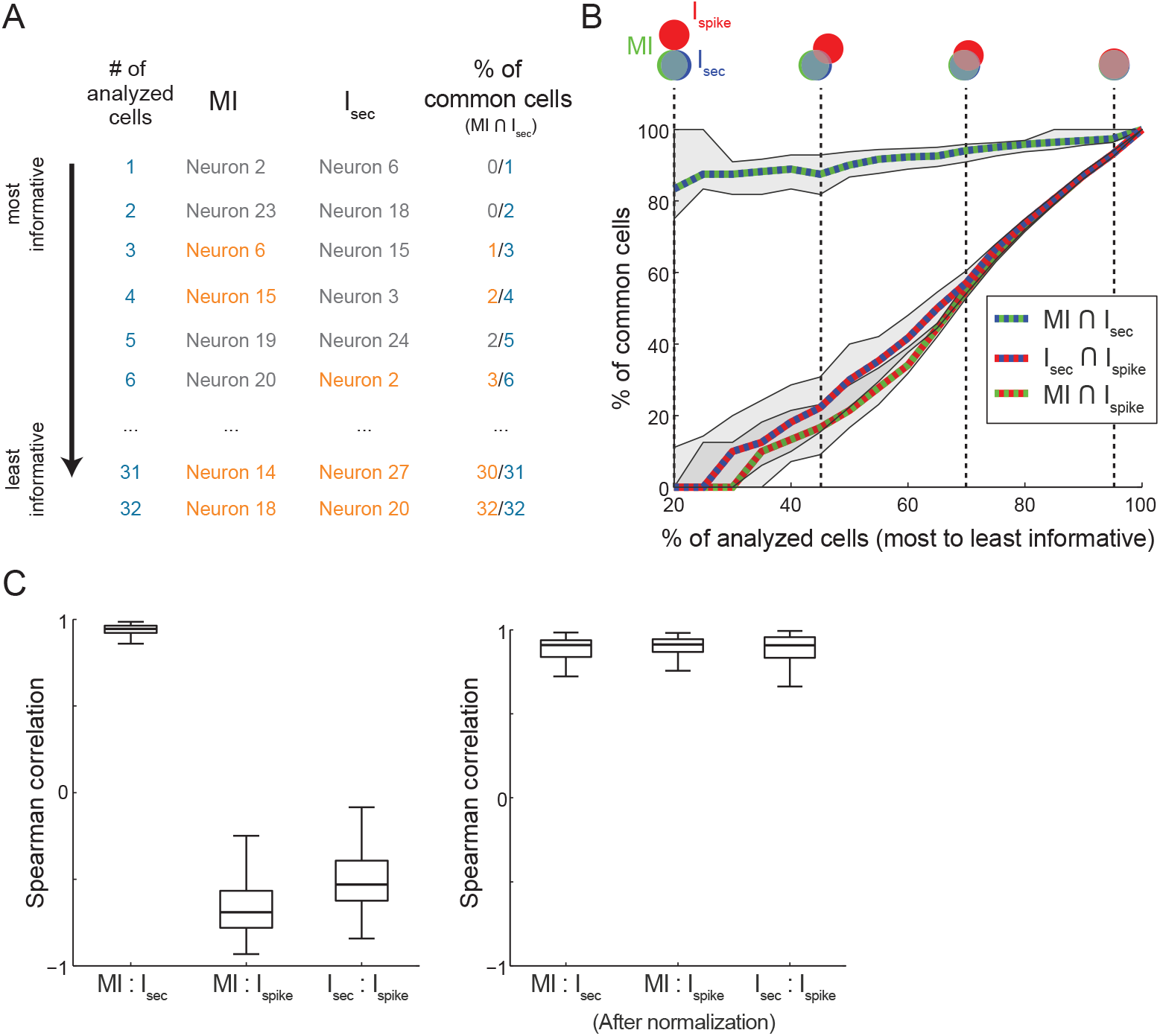
Comparison of the subpopulations of spatial cells according to each metric. **A.** For each linear track session, neurons were ranked from most to least informative according to I_sec_, I_spike_ or MI. The intersections (% of common cells) between the top informative neurons of each metric pair was computed considering the first N cells, with N varying from 1 to the total number of cells. The panel illustrates this procedure for a pair of metrics (MI and I_sec_) in one session. **B**. Median % of common cells plotted against the number of analyzed cells (in percentage from total cell number, N%) across all sessions for each pair of metrics. The shaded area represents interquartile range. Venn diagrams on top show the median intersection at the corresponding N%. Note the low intersection between I_sec_ and I_spike_ or MI and I_spike_. **C**. Boxplot distributions of the Spearman correlation between each pair of original (left) and normalized (right) metrics.

Finally, we investigated the relation between information and firing rate separately for putative pyramidal cells and interneurons. As shown in Figure 8A top panels, the range of information values for pyramidal cells and interneurons had greater overlap when estimated by MI, while the interneurons exhibited much lower information than pyramidal cells when estimated by I_sec_ and I_spike_. Moreover, in the case of I_spike_, the estimated information had a clear relation to firing rate. On the other hand, the normalized versions of each metric showed similar relations between firing rate and information (Figure 8A bottom panels), resembling the relation between firing rate and percentage of correct decoding (Figure 8B).

**Figure 8.**
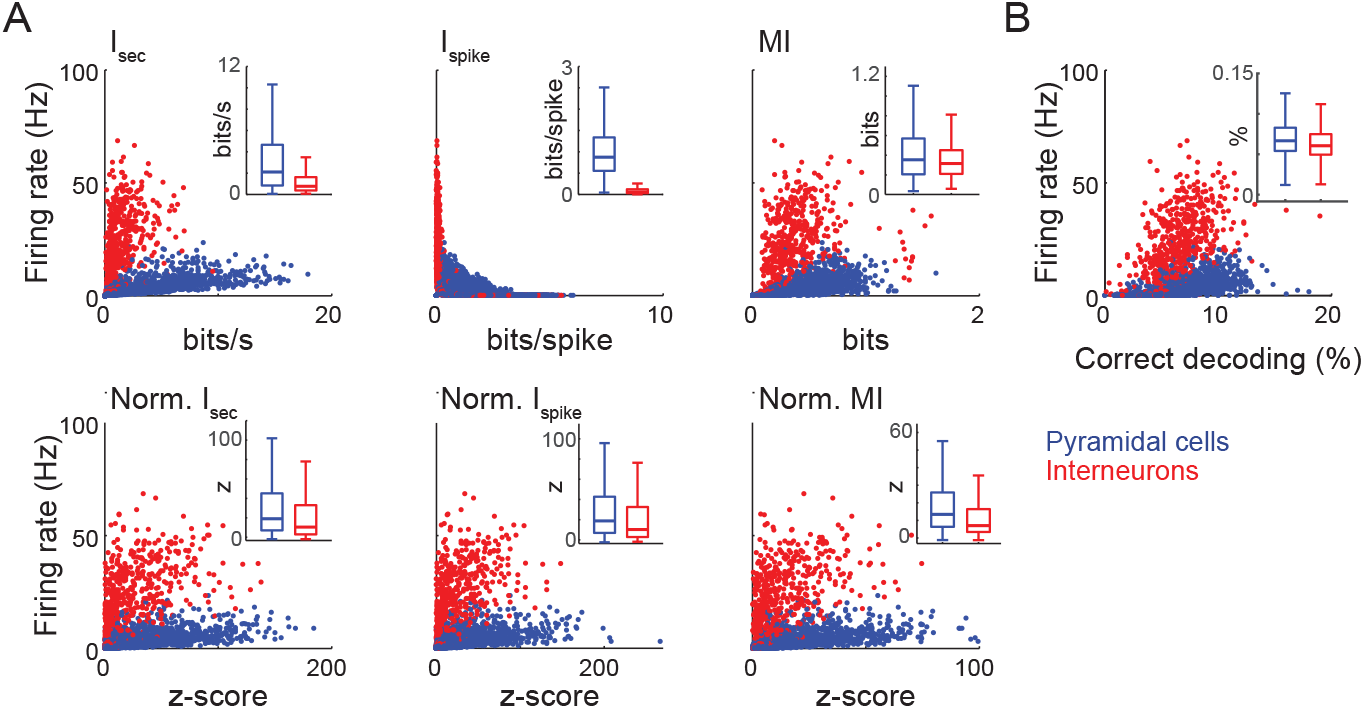
Information estimates for CA1 pyramidal cells and interneurons. **A.** Scatter plots of spatial information and firing rate for each metric (top) and its normalized version (bottom). Only neurons classified as pyramidal cells (blue) and interneurons (red) were considered. Insets show boxplot distributions of information values. **B**. Scatter plots of decoding performance and firing rate. Note inverse dependence of I_spike_ on firing rate, and greater overlap in the information range of interneurons and pyramidal cells for MI and the normalized metrics, akin to the overlap present in decoding performance.

## Discussion

We studied three metrics of spatial information using both simulated and real data. The performance of a Bayesian decoder was assumed to represent the gold standard of the true spatial information content of a cell. Decoders are used to infer the most likely stimulus that elicited a particular response; their performance is directly linked to the information about the stimulus contained in the response (Quiroga and Panzeri, 2009). In other words, if two variables are related, it might be possible to use one of them to decode the other. The information contained in the confusion matrix of a decoder provides a lower bound to the information between the two variables (Quiroga and Panzeri, 2009), allowing the use of decoding performance as an empirical estimate of the real (spatial) information of the cell (Robertson et al., 1999; Jensen and Lisman, 2000; Huxter et al., 2008; Lopes-dos-Santos et al., 2015). Under this framework, our simulations show that the mutual information (MI) better correlates with spatial decoding performance than I_sec_ and I_spike_. Similar results hold when running our analyses on real linear-track data (Figures 5 and 6), though I_sec_ had better correlation with correct decoding in real than simulated data. This might be explained by the absence of some types of spatial modulation considered in our simulated cells, along with the expected presence of classical place cells in CA1 recordings. Finally, we found that the correlation with decoding performance achieves the same levels among the metrics following a surrogate-based normalization.

The low correlation of I_sec_ and I_spike_ with correct decoding in simulated data is a consequence of the way these metrics quantify information. Because they use the average firing rate over trials, a single trial with high firing rate can bias the metrics towards higher information values. This issue was apparent for simulated cells that were spatially modulated only in some of the trials (cell types IV and V): cells with different number of modulated trials but same mean firing rate showed similar information values (trial consistency; see Figure 9). This contrasts with the intuitive notion that the more consistent the spatial modulation of a cell across trials, the higher its spatial information. Another characteristic of I_sec_ and I_spike_ metrics was their sensitivity to changes in basal firing rate. For instance, the same increase in firing rate but from different baseline levels (e.g., 0 to 5 Hz vs. 10 to 15 Hz) leads to different information values, favoring cells with low basal firing rate to have higher information (additive effect; Figure 9). Additionally, the I_sec_ metric is sensitive to changes in the mean firing rate of a cell upon a multiplicative factor (multiplicative effect; Figure 9).

**Figure 9:**
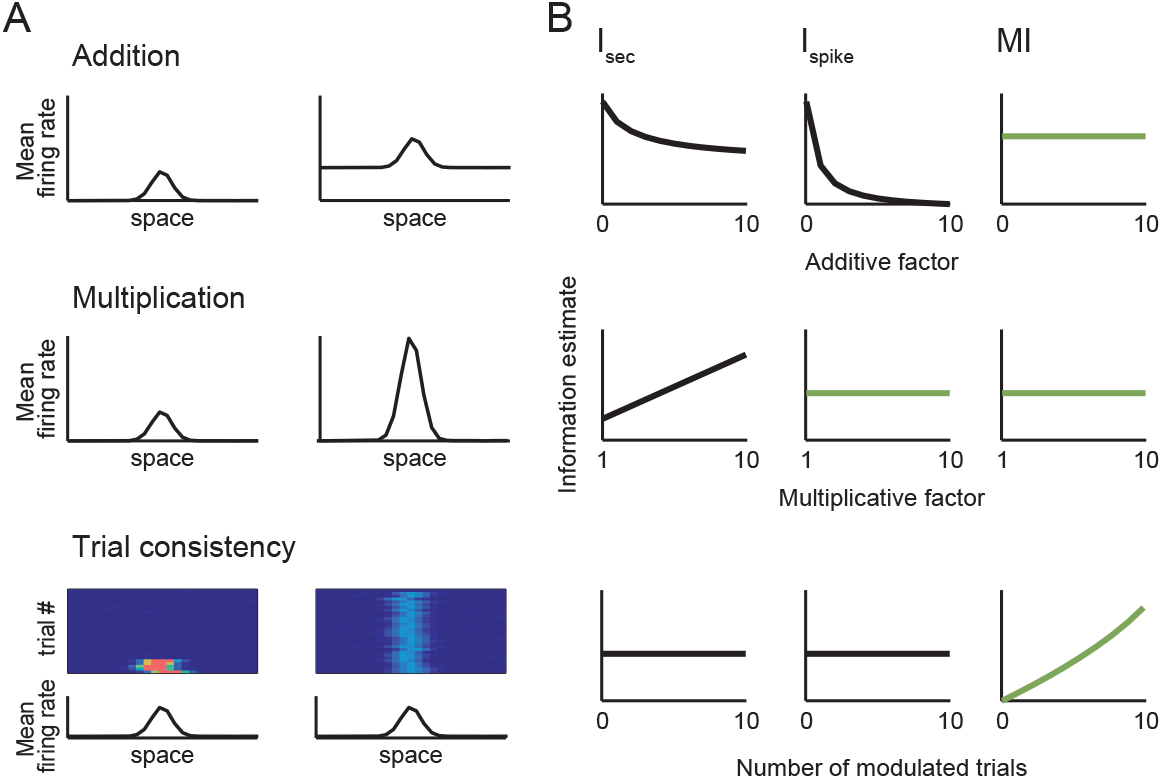
Influence of addition, multiplication and trial consistency on spatial metrics. **A.** Schematic examples of firing rate addition (top) and multiplication (middle), as well as of consistency over trials (bottom). In the latter case, the mean firing rate over trials was fixed. **B.** I_sec_, I_spike_ and MI for different parameters of the cases in A. I_sec_ and I_spike_ decay as the additive factor increases, while I_sec_ linearly increases with the multiplication factor. Only the MI increases with trial consistency. The green curves are cases in which the metric behaves similarly to the decoding performance.

Some of these effects may be due to the assumptions underlying the derivation of I_sec_ and I_spike_. Interpreting the information rate in bits per second may lead one to consider that this rate is valid for any amount of time. However, the estimated information rate might be valid only for small Δt, during which the cell emits only one or no spike. Since the information conveyed by a spike is not independent of the information from previous ones, the information rate may be overestimated as Δt increases (Skaggs et al., 1993). The amount of redundancy is then directly related to the firing rate of the cell, explaining why normalizing I_sec_ by the firing rate (i.e., I_spike_) impedes the overestimation caused by the multiplicative effect (Figure 9). The underestimation of the additive effect, on the other hand, is due to the fact that the firing rate of each spatial bin becomes closer to the average firing rate; that is, adding a constant to all firing rate bins reduces the variations across bins, making the ratio λ_i_ /λ closer to 1 and log(λ_i_ /λ) closer to 0. This effect is inherent to the definition of both I_sec_ and I_spike_, albeit the convergence to zero is faster for I_spike_ (Figure 9).

In contrast to I_sec_ and I_spike_, the MI was able to capture the true spatial information of the cells irrespective of rate-position map features. For instance, in the example cases shown in Figure 3, the MI was capable of properly estimating the information despite differences in mean firing rate (cell types I and II). This is because the MI is based on the probability of each firing rate value and not on the value itself. In other words, as opposed to I_sec_ and I_spike_, the MI is insensitive to additive and multiplicative effects (Figure 9). Moreover, the MI detected the increase of spatial information with higher trial consistency despite the constant mean firing rate across trials (cell type V). Notice that the MI takes into account the firing rate of every trial instead of the trial mean, making this metric more robust to inter-trial variability.

We introduced a normalization that estimates and corrects for the intrinsic bias present in the rate-position map. This bias is due to the fact that even shuffled firing rate maps will have information estimates above zero (i.e., random firing rate maps are seldom constant over space). After normalizing I_sec_ and I_spike_ by the mean and standard deviation of the shuffled distribution, we found an increase in their correlation with decoding performance, which reached the same level as the original MI (Figure 4). These normalizations may be useful under situations in which the definition of a trial is not possible (i.e., an open field) and thus the probabilities underlying the MI cannot be computed. It is worth noticing that binning the firing rate values to compute the MI might introduce information bias (Panzeri et al., 2007), which can also be corrected by the normalization.

These findings may have implications on what we know about spatial representations in the brain. For instance, previous studies measured spatial information in entorhinal cortex and hippocampus and found higher spatial information in putative excitatory than inhibitory neurons (Frank et al., 2000, 2001). However, this could reflect the bias of I_spike_ towards higher information values in cells with low mean firing rate (Figure 9B). Accordingly, we found that information estimates for putative pyramidal cells and interneurons in CA1 drastically differ for I_spike_ but not for MI (Figure 8). Moreover, our results further show that the three metrics can yield different subpopulations of spatially-modulated neurons (Figure 7). Surprisingly, despite their similar definition, we found very low intersection between neurons classified using I_spike_ and I_sec_ (Figure 7B).

More generally, our observations raise the question of what defines the spatial information of a neuron. For instance, most of the known correlates of space in the hippocampus focus on neurons that typically spike at very low rates outside their receptive fields (O’Keefe and Dostrovsky, 1971; Taube et al., 1990; Fyhn et al., 2004; Hafting et al., 2005). While I_spike_ and (mainly) I_sec_ work well to select cells with high spatial information in these cases, we wonder whether these metrics may have limited our understanding of spatial coding. In other words, the I_sec_ and I_spike_ are well suited to detect “canonical” place cells (i.e., cells whose spatial firing rate is a unimodal function centered on the place field), but may fail to detect other spatially informative neurons whose firing rate map is not that of a canonical place cell. Notice that adding a constant factor to a spatial firing rate map does not influence decoding performance nor the information estimated by the original MI, but decreases the amount of spatial information estimated by I_sec_ and I_spike_ (Figure 9). While some researchers may intuitively consider that cells which are silent outside the place field convey more information when they spike than cells with high basal firing rates, others may be more concerned as to whether it is possible or not to decode the animal position from the firing rate of the cell. The latter would further argue that every cell that carries information about the animal’s position (as retrieved by a decoder) could be called a “place cell”, independently of place field shape or mean firing rate. Noteworthy, the existence of unconventional spatial representations has been recently demonstrated for medial entorhinal cortex neurons (Diehl et al., 2017; Hardcastle et al., 2017). In any case, our results suggest that the MI is a suitable metric to capture other types of less canonical spatial correlates that may have gone undetected so far. In cases where the MI cannot be computed, the normalization of I_sec_ and I_spike_ may constitute good alternatives.

## Acknowledgements

This work was supported by CAPES and CNPq, Brazil. We thank the Buzsáki laboratory for making data publicly available at http://crcns.org/, a data-sharing website supported by NSF and NIH, USA. We thank D. Laplagne and R. Romcy-Pereira for critical discussions.

## References

Diehl, G.W., Hon, O.J., Leutgeb, S., Leutgeb, J.K., 2017. Grid and nongrid cells in medial entorhinal cortex represent spatial location and environmental features with complementary coding schemes. Neuron 94, 83–92.

Ego-Stengel, V., Wilson, M.A., 2007. Spatial selectivity and theta phase precession in CA1 interneurons. Hippocampus 17, 161–174.

Eichenbaum, H., 2000. A cortical–hippocampal system for declarative memory. Nat. Rev. Neurosci. 1, 41–50.

Frank, L.M., Brown, E.N., Wilson, M., 2000. Trajectory encoding in the hippocampus and entorhinal cortex. Neuron 27, 169–178.

Frank, L.M., Brown, E.N., Wilson, M.A., 2001. A comparison of the firing properties of putative excitatory and inhibitory neurons from CA1 and the entorhinal cortex. J. Neurophysiol. 86, 2029–2040.

Fyhn, M., Molden, S., Witter, M.P., Moser, E.I., Moser, M.-B., 2004. Spatial representation in the entorhinal cortex. Science 305, 1258–1264.

Hafting, T., Fyhn, M., Molden, S., Moser, M.-B., Moser, E.I., 2005. Microstructure of a spatial map in the entorhinal cortex. Nature 436, 801–806.

Hardcastle, K., Maheswaranathan, N., Ganguli, S., Giocomo, L.M., 2017. A multiplexed, heterogeneous, and adaptive code for navigation in medial entorhinal cortex. Neuron 94, 375–387.

Huxter, J.R., Senior, T.J., Allen, K., Csicsvari, J., 2008. Theta phase-specific codes for two-dimensional position, trajectory and heading in the hippocampus. Nat. Neurosci. 11, 587–594.

Jensen, O., Lisman, J.E., 2000. Position reconstruction from an ensemble of hippocampal place cells: contribution of theta phase coding. J. Neurophysiol. 83, 2602–2609.

John, G.H., Langley, P., 1995. Estimating continuous distributions in Bayesian classifiers. In: Proceedings of the Eleventh Conference on Uncertainty in Artificial Intelligence, pp. 338–345. Morgan Kaufmann Publishers Inc.

Kropff, E., Carmichael, J.E., Moser, M.-B., Moser, E.I., 2015. Speed cells in the medial entorhinal cortex. Nature 523, 419.

Lopes-dos-Santos, V., Panzeri, S., Kayser, C., Diamond, M.E., Quiroga, R.Q., 2015. Extracting information in spike time patterns with wavelets and information theory. J. Neurophysiol. 113, 1015–1033.

Mizuseki, K., Sirota, A., Pastalkova, E., Buzsáki, G., 2009. Theta oscillations provide temporal windows for local circuit computation in the entorhinal-hippocampal loop. Neuron, 64, 267–280.

Mizuseki, K., Sirota, A., Pastalkova, E., Diba, K., Buzsáki, G., 2013. Multiple single unit recordings from different rat hippocampal and entorhinal regions while the animals were performing multiple behavioral tasks. CRCNS Org.

Morris, R.G.M., Garrud, P., Rawlins, J.N.P., O’Keefe, J., 1982. Place navigation impaired in rats with hippocampal lesions. Nature 297, 681–683.

Moser, E.I., Kropff, E., Moser, M.-B., 2008. Place cells, grid cells, and the brain’s spatial representation system. Annu. Rev. Neurosci. 31, 69–89.

O’Keefe, J., Dostrovsky, J., 1971. The hippocampus as a spatial map. Preliminary evidence from unit activity in the freely-moving rat. Brain Res. 34, 171–175.

O’Keefe, J., Recce, M.L., 1993. Phase relationship between hippocampal place units and the EEG theta rhythm. Hippocampus 3, 317–330.

Panzeri, S., Senatore, R., Montemurro, M.A., Petersen, R.S., 2007. Correcting for the sampling bias problem in spike train information measures. J. Neurophysiol. 98, 1064–1072.

Quiroga, R.Q., Panzeri, S., 2009. Extracting information from neuronal populations: information theory and decoding approaches. Nat. Rev. Neurosci. 10, 173–185.

Robertson, R.G., Rolls, E.T., Georges-François, P., Panzeri, S., others, 1999. Head direction cells in the primate pre-subiculum. Hippocampus 9, 206–219.

Scoville, W.B., Milner, B., 1957. Loss of recent memory after bilateral hippocampal lesions. J. Neurol. Neurosurg. Psychiatry 20, 11.

Shannon, C.E., 1948. A mathematical theory of communication, Part I, Part II. Bell Syst Tech J 27, 623–656.

Skaggs, W.E., McNaughton, B.L., Gothard, K.M., Markus, E.J., 1993. An information-theoretic approach to deciphering the hippocampal code, in: Advances in Neural Information Processing Systems 5. eds. Hanson, S.J., Giles, C.L. & Cowan, J.D., pp. 1030–1037.

Skaggs, W.E., McNaughton, B.L., Wilson, M.A., Barnes, C.A., 1996. Theta phase precession in hippocampal neuronal populations and the compression of temporal sequences. Hippocampus 6, 149–172.

Taube, J.S., Muller, R.U., Ranck, J.B., 1990. Head-direction cells recorded from the postsubiculum in freely moving rats. I. Description and quantitative analysis. J. Neurosci. 10, 420–435.

Zola-Morgan, S.M., Squire, L.R., 1990. The primate hippocampal formation: evidence for a time-limited role in memory storage. Science 250, 288–290.

